# Cryo-EM structure of Alzheimer’s disease tau filaments with PET ligand MK-6240

**DOI:** 10.1101/2023.09.22.558671

**Authors:** Peter Kunach, Jaime Vaquer-Alicea, Matthew S. Smith, Robert Hopewell, Jim Monistrol, Luc Moquin, Joseph Therriault, Cecile Tissot, Nesrine Rahmouni, Gassan Massarweh, Jean-Paul Soucy, Marie-Christine Guiot, Brian K. Shoichet, Pedro Rosa-Neto, Marc I. Diamond, Sarah H. Shahmoradian

## Abstract

Positron Emission Tomography (PET) ligands have advanced Alzheimer’s disease (AD) diagnosis and treatment. Using autoradiography and cryo-EM, we identified AD brain tissue with elevated tau burden, purified filaments, and determined the structure of second-generation high avidity PET ligand MK-6240 at 2.31 Å resolution, which bound at a 1:1 ratio within the cleft of tau paired-helical filament (PHF), engaging with glutamine 351, lysine K353, and isoleucine 360. This information elucidates the basis of MK-6240 PET in quantifying PHF deposits in AD and may facilitate the structure-based design of superior ligands against tau amyloids.

## Main

Alzheimer’s Disease (AD) is characterized by the progressive accumulation of amyloid-beta (Aβ) plaques and neurofibrillary tangles (NFTs), with tau protein aggregates forming paired helical filaments (PHFs) and straight filaments (SFs) central to AD pathogenesis (1-6). Positron emission tomography (PET) imaging with Aβ and tau ligands has enhanced diagnostic accuracy and understanding of AD progression (7). First-generation tau-PET ligands have enabled in vivo detection of tau tangles and have predictive capabilities for brain atrophy and cognitive decline in pre-symptomatic individuals (8-14).

Building on this, second-generation tau-PET ligands have been developed to improve specificity, pharmacokinetics, and avidity. These ligands are based on optimized chemical structures such as pyridoindole, phenyl-butadienyl-benzothiazoles, and quinoline/benzimidazole derivatives (15-17). Specifically, pyrrolopyridinyl isoquinoline amine derivatives like [18F]MK-6240 have shown superior binding to tau tangles compared to first-generation tracers (15). The development of second-generation tau-PET ligands, such as [18F]MK-6240, is vital for early AD detection, disease staging, and therapeutic intervention evaluation (18-23).

While ligand development methods have been productive (24-28), cryogenic electron microscopy (cryo-EM) offers atomic-resolution insights into filament binding (29-31). Our study focuses on MK-6240, which is increasingly used in clinical studies (23), and determined its structure in complex with AD-derived tau fibrils using cryo-EM.

We started with a neuropathologically confirmed AD case, scored as A3, B3, C3 according to Thal phases (Aβ), Braak stage (NFT), and CERAD score (Neuritic plaque). We selected frontal cortical tissue to purify AD filaments (Extended Data Fig. 1). Guided by autoradiography with [18F]MK-6240, we dissected regions with high AD tau filament concentration (Extended Data Fig. 2A) and extracted sarkosyl-insoluble tau filaments using standard methods (Extended Data Fig. 2B) (32). Cryo-EM confirmed the presence of isolated tau filaments (Extended Data Fig. 3A/B), and in vivo binding of the purified tau filaments to MK-6240 was validated by inoculation into the rat hippocampus (Extended Data Fig. 4 and 5).

Prior to cryo-EM imaging, we incubated tau PHFs from the frontal cortex with 20 µM MK-6240 (Fig. 1A) or 3% DMSO as a control (Extended Data Fig. 6A). Cryo-EM reconstructions using RELION revealed PHF conformations consistent with known structures, with an estimated resolution of 3.0 Å (Fourier Shell Correlation, FSC = 0.143) (Extended Data Fig. 7). Analysis of PHFs complexed with MK-6240 yielded a higher resolution of 2.31 Å (FSC = 0.143) (Extended Data Fig. 8). The density corresponding to MK-6240 was discernible at 12.5 standard deviations above the noise, and the amino acid main chain was extended up to 16 standard deviations (Fig. 1B). Notably, MK-6240 molecules exhibited a 1:1 stoichiometry with tau monomer rungs (Fig. 1C/G), a pattern that was mirrored in the cross-sections (Extended Data Fig. 6A), presented at a sigma contrast value of 7.5.

**Figure. 1:**
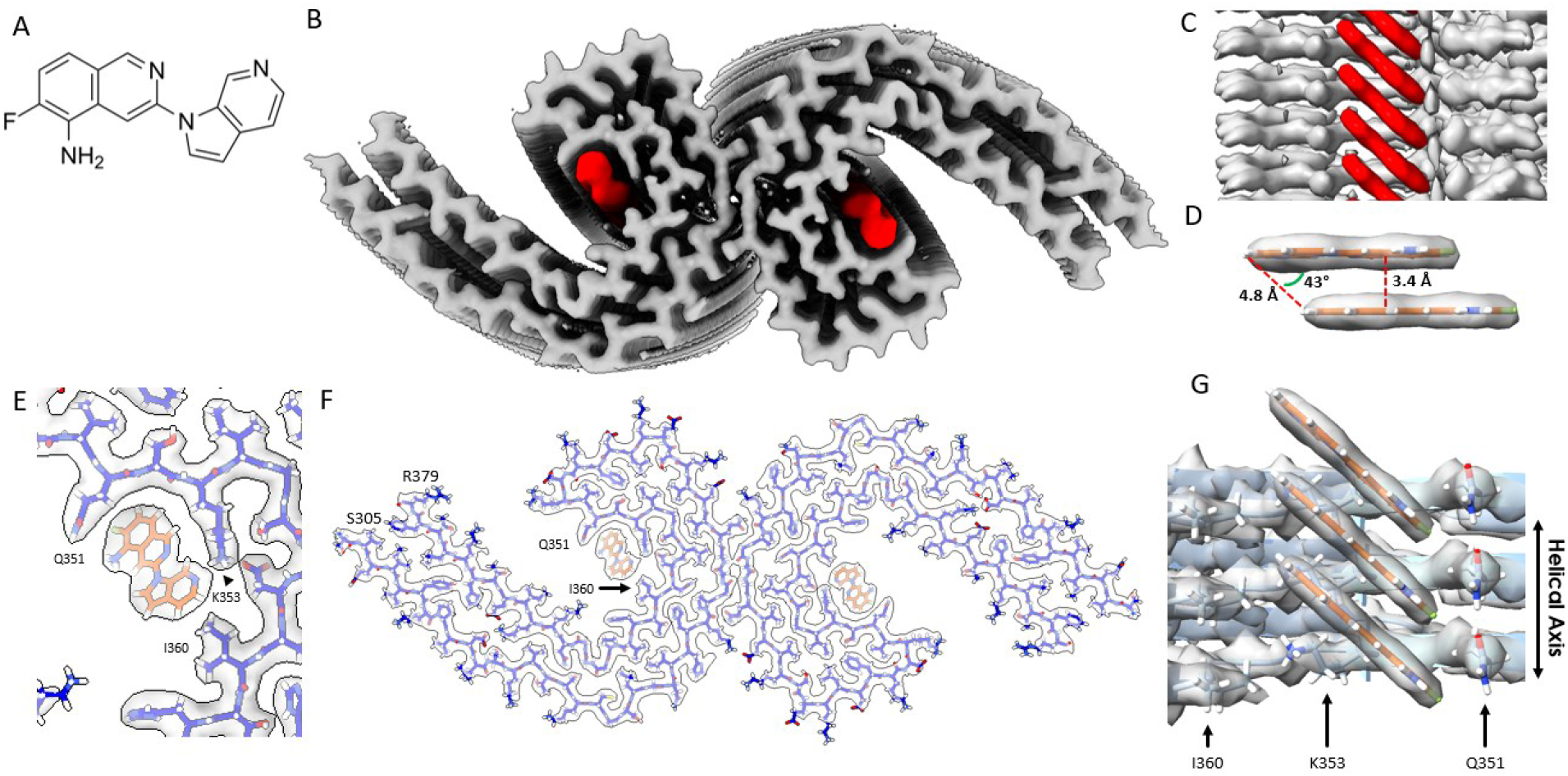
Structure of MK-6240 bound to tau PHF. **(A)** Molecular structure of MK-6240. **(B)** Cryo-EM map of AD tau PHFs (Grey) with bound MK-6240 (Red). **(C)** Side-view depiction of MK-6240 (red) situated within the binding pocket of the cleft within the AD tau PHF (Grey). MK-6240 adopted a stacked arrangement perpendicular to the fibril axis. One MK-6240 molecule spanned approximately two tau monomer rungs. **(D)** Isolated atomic representation of MK-6240 with its binding orientation. Peripheral atoms (alpha-carbon Hydrogen, left; and Fluorine, right) exhibit a 4.8 Å distance between them. Between heterocycles or parallel atoms within MK-6240 molecules, a ~3.3 Å distance is observed. **(E)** Close-up depiction of the binding site, accentuating the proximity of the three amino acids (Q351, K353, and I360) to MK-6240. **(F)** Atomic model showcasing the MK-6240 binding site on paired helical filaments. Cryo-EM density (white) is juxtaposed with the atomic model of the tau fold (blue) and MK-6240 (orange). **(G)** Side-view, zoomed-in perspective (from within the cleft) of the MK-6240 binding site, revealing the ~1:1 stoichiometry and the angle that aligns individual MK-6240 molecules with both themselves and the filament axis.

Using Chimera, we fit the atomic coordinates of MK-6240 within the largest non-protein density in the C-shaped cavity of each tau protofilament, predicting packing interactions with amino acids Q351 and I360, and a hydrogen-bond with K353 (Fig. 1 E/F/G). The alignment of the 6-Azaindole ring of MK-6240 near K353 was stabilized by an inter-residue ion pair with neighboring D358. Hydrophobic contacts were formed between the fluorine (Fl) of the isoquinoline ring and adjacent carbons with Q351’s aliphatic chain, while the Fl was oriented alongside Q351’s sidechain carboxamide. This positioning facilitated van der Waals interactions with I360 and the 6-Azaindole ring, leaving the Fl and primary amine groups solvent exposed and supported by electrostatic complementarity with polar residues (Fig. 2A).

**Figure. 2:**
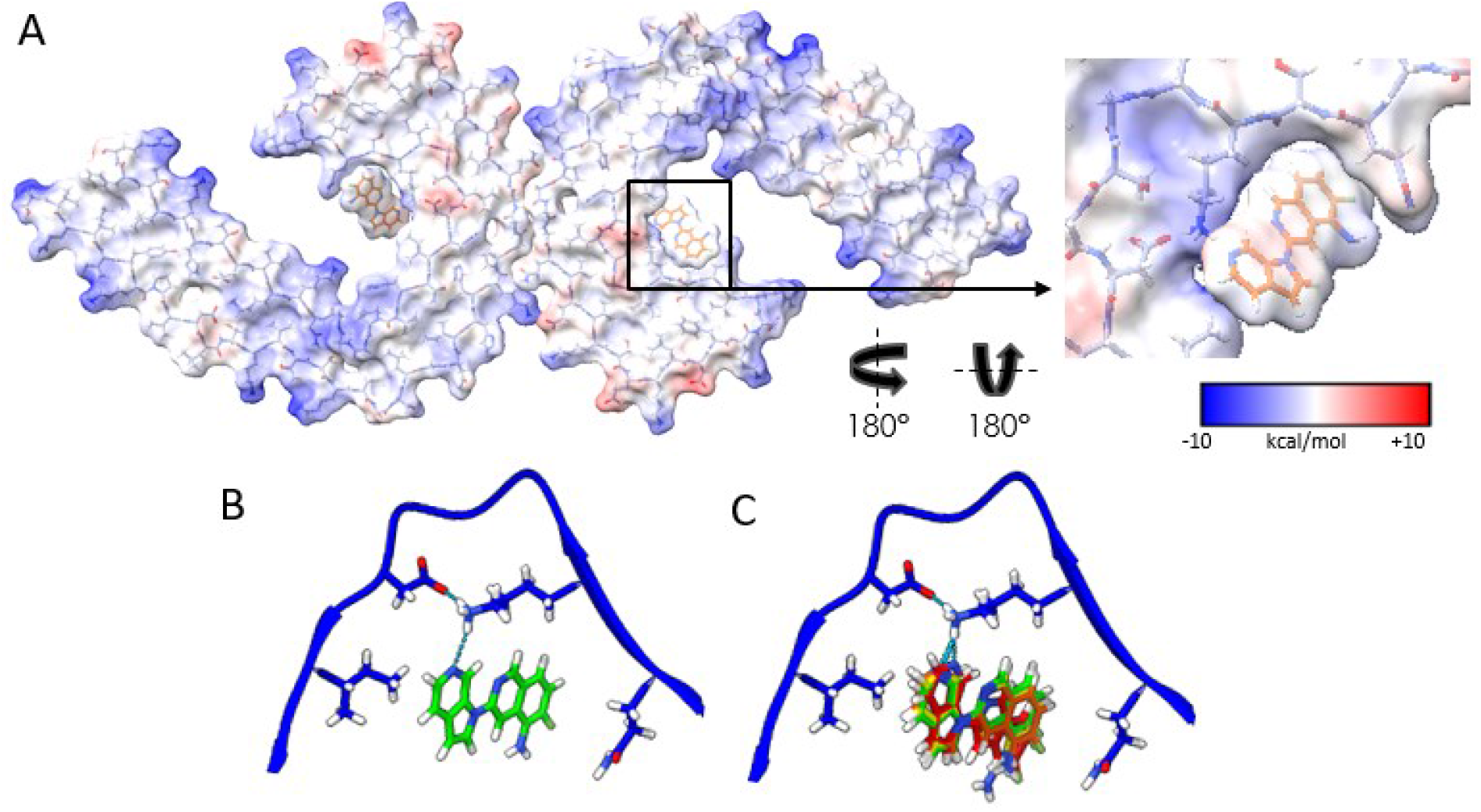
Electrostatic interactions between MK-6240 and PHFs and correspondence of the docking prediction to the experimental result. **(A)** Coulombic energy map depicting the cryo-EM density of the binding pocket, with amino acids from the tau fold and MK-6240. **(B)** Optimal pose resulting from MK-6240 binding via symmetry docking, revealing hydrogen bonding involving the secondary amine of the azaindole ring and K353. Minimized energetic scores align with the manually docked pose of MK-6240 in the cryo-EM density as shown in (B). **(C)** The top four poses (Green > Yellow > Orange > Red) generated through symmetry docking, showcasing robust alignment and a predilection for the orientation of the secondary amine of the azaindole ring system to facilitate interaction with K353. Flexibility variations toward the solvent-exposed region are observed among these poses.

MK-6240’s binding configuration was characterized by a slanted columnar arrangement from staggered alignment, revealing a 4.8 Å spacing at equivalent distal atoms (Fig. 1D), similar to the separation between AD PHF protofilament rungs (32). This alignment avoided steric hindrance from MK-6240’s halogen, with a 3.4 Å distance between the closest atoms and overlapping aromatic rings (Fig. 1D) suitable for an offset stacking arrangement (46). The columnated MK-6240, inclined at approximately 43°, conformed to the fibril’s twist, ensuring stable binding and mirroring cryo-EM structures of tau with GTP-1. This orientation prevented collisions between projecting MK-6240 molecules and tau filament, possibly involving hydrogen bonding between the azaindole ring’s amine and K353 (Fig. 2A/B) and inducing torsion on the mainchain between K340 and G355 (Extended Data Fig. 6C).

In addition to the unambiguous binding site (red arrow), we observed an additional density, potentially indicating a substoichiometric interaction involving I360 and H362 (orange arrow) (Extended Data Fig. 6A). This density was absent in our control structure. We predict that MK-6240 could fold a salt bridge with H362 in a non-columnated binding mode. These findings, along with the reported subtle densities in other PHF maps, require further investigation to determine their relevance.

To further understand MK-6240-induced side chain rearrangement, we overlaid the difference map generated from the PHF+MK-6240 and PHF maps (Salmon density, Extended Data Fig. 6B) onto the unbound PHF map (Grey density, Extended Data Fig. 6B). After aligning the structures with the remaining amino acids (i.e., 306-339 and 356-374), we compared the model from the PHF+MK-6240 map to the unbound map (Extended Data Fig. 6C). The α-carbon root-mean-square deviations (RMSDs) of residues 340-355 from the PHF+MK-6240 structure were 1.1, 1.6, and 1.2 Å when compared to published unbound PHF models 5o3l, 6HRE, and the control structure. The α-carbon RMSD between the control structure and unbound PHF models 5o3l and 6HRE was 0.65 Å each, indicating highly similar unbound structures (Extended Data Fig. 9A). Additionally, we assessed values from Merz et al., (31) revealing α-carbon RMSD of residues 340-355 between the PHF+GTP-1, and the unbound PHF and PHF+MK-6240 models, with RMSD values of 0.56 and 1.65 Å, respectively (Extended Data Fig. 9B). These data suggest that the protein backbone and side chains are flexible enough to accommodate MK-6240 binding, akin to the binding of GTP-1.

It has been well documented that the PHF is the predominant tau polymorph in Alzheimer’s disease (4,5), however, it has also been observed in various other neurodegenerative conditions, such as Familial British Dementia, Familial Danish Dementia, and PrP Cerebral Amyloid Angiopathy (6). Our data suggests that MK-6240 is a useful tool for measuring PHFs. Therefore, we predict that the utility of MK-6240 in vivo is contingent on the prevalence of the PHF polymorph in the respective condition.

To inform our structural model, we used a new implementation of UCSF DOCK 3.8. SymDOCK (Manuscript in preparation, Matthew S. Smith, et al., Brian K Shoichet), checks that prospective poses can form a stack using the symmetry of the protein and evaluates the ligand-ligand van der Waals energy. We docked MK-6240 against the structure of GTP-1 bound to AD PHF tau (31). Among the top 50 poses (by DOCK score) generated for MK-6240, the 9th-best one by docking score fits the electron density best (Figure 2C), though the top four docking poses also closely resemble the experimental structure (Figure 2D).

It is still unknown whether the interaction of PET ligands with tau filaments, as revealed by cryo-EM structures, accurately reflects their binding modes *in vivo* (30). For example, inaccurate structural models could be derived as a result of saturating ligand concentrations used in cryo-EM experiments. Future work will aim to resolve structures using sub-stoichiometric conditions.

While stacked interactions between ligands are rare in traditional protein-ligand complexes, they are frequently observed in fibril-ligand complexes (30, 31, 45). It seems that ligand-ligand interactions act as a stabilizing molecular glue. To this end, MK-6240 has little interactions with its target protein than is typical for such high-affinity interactions. The surface area involved in ligand-ligand interactions (243 A^2^) is greater than that in the ligand-protein interface (208 A^2^), resulting in 46% of the solvent accessible surface area (SASA) of an MK-6240 monomer alone being buried by the protein, or 69% of the SASA of an MK-6240 monomer within the protein and a stack of MK-6240 molecules. These values are similar to what has been previously calculated for GTP-1 bound to tau fibrils (30).

Liu et al.’s (48) highlighted the importance of binding modes in determining ligand specificity. Structural analyses show varying binding modes among tau-PET ligands and amyloid disaggregants: APN-1607 binds in a parallel fashion, ECGC adopts a flat stacking mode, while MK-6240 and GTP-1 both use diagonal stacking. Future experiments will help clarify how binding mode contributes to specificity.

Our study represents the first examination of the molecular binding interface of PHFs derived from AD brain, complexed with MK-6240, a second-generation, high avidity tau-PET ligand. The binding mode involves specific amino acid contacts and a stacked arrangement with pi-pi aromatic interaction, reflecting a 1:1 stoichiometric ratio. These findings will guide rational PET ligand design for PHFs and other amyloid targets, contributing to the advancement of diagnostic tools in neurodegenerative diseases. Future research should investigate the additional density potentially indicating a substoichiometric interaction and the unknown ability of the SF mainchain to accommodate MK-6240 binding.

## Methods

### Clinical History and Neuropathology

We extracted tau filaments from the frontal cortex of a neuropathologically confirmed case of AD, an 87-year-old female native of Quebec, Canada. She had developed severe cognitive and functional symptoms in the 8 years prior to death. MRI indicated severe subcortical atherosclerosis-associated leuco-encephalopathy.

Neuropathological ABC categorization was: A3, B3, C3. Additional pathological findings included cerebral amyloid angiopathy, vascular pathology related to hypertension and/or atherosclerosis, and severe diabetes. The dentate gyrus and the CA1 region of the hippocampus, which had limited TDP-43 co-pathology, were excluded from the selection process. We observed no evidence of Lewy body pathology. The patient exhibited severe amnestic symptoms starting 8 years prior to death, experiencing difficulties in word-finding, and expressing coherent ideas. Assistance was required for eating, maintaining personal hygiene, and climbing stairs. Impairments in temporal and spatial orientation were noted, although recognition of familiar individuals remained intact. MRI results indicated severe subcortical atherosclerosis-associated leuco-encephalopathy. Over time, the patient’s agitation escalated, leading to conflicts with fellow residents, involving verbal insults, physical altercations, and vocalizing animal sounds. Progressive cognitive decline was evident through decreasing MMSE scores: 20/30 (2004), 13/30 (2008), 12/30 (March 2009), 12/30 (September 2009), 10/30 (2010), 13/30 (April 2011), and 1/30 (July 2011). The patient passed away in 2012.

For detailed neuropathological characterization, please refer to Extended Data Fig. 1.

### Immunohistochemistry

Immunohistochemistry was performed on formalin fixed paraffin embedded (FFPE) 5-micron sections, using a Ventana Discovery Ultra automatic stainer. The Ventana staining protocol included a 24 min pretreatment with Cell Conditioner 1 (CC1 Ventana), followed by incubation with the primary antibody for 60 min. Antibody was diluted in Ventana dilution buffer as follows: AT8 (phosphor-Tau Ser202, Thr205) (InVitrogen) 1/1000. Detection was performed using Discovery OmniMap DAB anti Ms RUO. (Ventana). The slides were scanned at 20X on an Aperio ScanScope XP and visualized using AperioImageScope software.

### Tissue Guided Autoradiography

Guided by autoradiography with [^18^F]MK-6240, tau filament-rich regions were recovered. Flash-frozen tissues were sectioned into 20 µm thick slices using a freezing sliding microtome (Leica CM3050 S) at − 15 °C. The sections were then placed on coated microscope slides and allowed to thaw at room temperature. Following this, the samples were air-dried and preincubated with a phosphate-buffered saline (PBS) solution (pH 7.4) containing 1% bovine serum albumin for 10 minutes to remove endogenous ligands. After additional air-drying, the samples were incubated with 20.4 MBq of [^18^F]MK-6240 in 600 mL of PBS for 150 minutes. Next, the tissues were dipped three times in PBS, followed by a dip in 4°C distilled H_2_O and dried under a stream of cool air. Finally, the samples were exposed to a radioluminographic imaging plate (Fujifilm BA SMS2025) for 20 minutes, and the activity quantified as photostimulated luminescence units per mm^2^ using ImageGauge software 4.0 (Fujifilm Medical Systems, Inc.).

The cortical regions utilized in the study are highlighted in Extended Data Fig. 2.

### Extraction of Tau Filaments

Sarkosyl-insoluble material was extracted from the cortical grey matter. Tissues were homogenized using a protocol adapted from the preprint by Fitzpatrick *et al*., (32). A total of 4 g of tissue was homogenized in extraction buffer (10 mM Tris-HCl, pH 7.4, 10% Sucrose, 0.8 M NaCl, 5 mM EDTA, 1 mM EGTA, and a protease phosphatase inhibitor (Roche) at a ratio of 20 mL/g. The homogenates were centrifuged at 3,900g for 10 minutes, followed by incubation with 2% sarkosyl for 1 hour at room temperature with constant stirring. The resulting supernatant was then spun at 100,000g for 1 hour at 4°C to obtain sarkosyl-insoluble pellets. These pellets were resuspended in extraction buffer at a ratio of 1 mL/g of starting material and centrifuged at 3,000g for 5 minutes at 4°C. The resulting supernatant was diluted 5-fold in a buffer containing 50 mM Tris-HCl, pH 7.5, 0.15 M NaCl, 10% Sucrose, and 0.2% sarkosyl, and then subjected to centrifugation at 100,000g for 30 minutes at 4°C. The pellet obtained was resuspended in Tau Buffer (20 mM Tris-HCl, pH 7.4, 100 mM NaCl) at a ratio of 250 µL/g of starting material and centrifuged at 100,000g for 30 minutes. Finally, the resulting pellet was resuspended in Tau Buffer at a ratio of 75 µL/g of starting material.

### Incubation of AD-tau filaments with MK-6240

MK-6240 was purchased from NucMedCor. 10 mg of MK-6240 was dissolved in 1 mL of 100% DMSO and then titrated to a working solution 600 µM. Certificate of Analysis included at the end of this manuscript. 1 µL of working solution was added to 30 µL of sarkosyl-insoluble fraction from AD brain for a final concentration of 20 µM MK-6240 and 3% DMSO and incubated at room temperature for 3 hours.

### Cryo-EM Grid Preparation

To ensure saturation of all potential binding sites, the sarkosyl-insoluble fraction was incubated with MK-6240 at a concentration of 20 µM prior to vitrification. Subsequently, the sample was centrifuged at 2,500 x g for 1 min. For cryo-electron microscopy, a volume of 3.5 µL of the sample was applied to a glow-discharged holey carbon grid (Quantifoil Cu R1.2/1.3, 300 mesh) and rapidly frozen in liquid ethane using an FEI Vitrobot Mark IV (Thermo Fischer Scientific). The grids were then loaded onto a 300 kV Titan Krios microscope for imaging. Movies consisting of 60 frames were acquired using a Gatan K3 summit detector in CDS mode. The total electron dose administered was 62 e-/Å2, and the super resolution pixel size was set at 0.415Å. To eliminate inelastically scattered electrons, a GIF quantum energy filter (Gatan) with a slit width of 20 eV was employed. Further details can be found in Extended Data Table 1.

### Helical Reconstruction

The collected movies were binned by two, motion corrected, and dose weighted using the motion correction implementation provided by RELION 4.0 (37). Aligned and dose-weighted micrographs were then used to estimate the contrast transfer function (CTF) using CTFFIND-4.1 (38). The image processing steps were guided by the protocols described by Sofia Lovestam et al., 2022 (39), and Sjors Scheres, 2020 (40).

Initially, filaments were manually picked and extracted using a box size of approximately 850 Å (1024 pixels) with an inter-box distance of 14.25 Å (three subunits), which was then downscaled by a factor of four. A first round of reference-free 2D classification was performed using a mask diameter of 840 Å and a tube diameter of 200 Å, resulting in predominantly PHF 2D-class averages. The highest resolution averages were selected, and particles were re-extracted using a box size of 280 Å without rescaling. The best 2D-class averages were chosen.

For de novo 3D initial model generation, the relion_helix_inimodel2d program (40) was employed with a cross-over distance of 740 Å. A first 3D auto-refinement was conducted using the initial model as a reference. Subsequently, a second 3D auto-refinement was performed using a soft-edged solvent-flattened mask (lowpass filtered, 15 Å) generated from the output of the first refinement. Finally, a pseudo 2-1 helical screw symmetry was imposed (twist: 179.42°, rise: 2.37 Å), and a search range for helical rise and twist was set to accurately refine the half-maps.

Bayesian polishing was performed on the particles, and the polished particles were used for a fourth round of 3D auto-refinement. Three rounds of CTF-refinement were subsequently conducted. Following this, a 3D classification without further image alignment was performed to eliminate segments that contributed to suboptimal 3D reconstructions. For further refinement, a single class containing 32,986 particles was selected. These particles underwent another set of auto-refinement, particle polishing, and three rounds of CTF refinements to generate a final reconstruction. Masking and post-processing techniques were applied, resulting in a map with a resolution of 2.31 Å using an FSC criterion of 0.143. The reconstruction incorporated pseudo 2-1 helical screw symmetry with a twist of 179.462° and a rise of 2.382 Å using the relion_helix_toolbox program (40), with a central Z length of 30%.

The post-processed map obtained from the above steps was then utilized for model building and refinement. Additional details can be found in Extended Data Fig. 7/8 and Extended Data Table 1.

### Model Building and Refinement

A model consisting of three rungs was initially generated using model_angelo (41), specifically utilizing the sequence S305-R379 from the microtubule binding domain. An individual rung was then extracted from the model using Chimera (42) and subjected to real-space refinement with ISolde (43). Model validation was performed using Phenix (44). Additional information and details can be found in Extended Data Table 1.

We translated and rotated a copy of the 3 Rung model refined from our cryo-EM map into a model containing 6 tau rungs because a single ligand interacts with 4 rungs. Running this calculation for the 3-rung model would have left some of the middle ligand hanging off the side of the modeled protein. Doing so undervalues the buried SASA, so we calculated these values for the 6-rung model. We calculated the buried SASA for an MK-6240 monomer in the protein fibril using Chimera. See extended data for calculations.

### Animals

Five 3-month-old male Sprague Dawley rats were used. They were housed at the Douglas Mental Health University Institute animal facility on a 12/12 hour light/dark cycle and had *ad libitum* access to food and water. All procedures described were performed in accordance with the Canadian Council on Animal Care guidelines and approved by the McGill University Animal Ethics Committee.

### Surgery

Rats were administered 0.1cc medetomidine and deeply anesthetized with 5% isoflurane at 1 L/min oxygen. Upon reaching a lack of withdrawal reflex, isoflurane was adjusted to approximately 1% for maintenance. The rat was immobilized in a stereotaxic frame (David Kopf Instruments), with drill and microinjection pump attachments. Animals were aseptically inoculated with human brain extracts in the dorsal hippocampus (bregma: −2 mm; lateral: - 3.5 mm; depth: −3.5mm) using a 5 µL Hamilton syringe. Injection needles were lowered to - 3.75mm for approximately 1 min to create a cavity at the injection site, followed by a return to -3.5mm where the inoculum was delivered. Each of the two injection sites received 1.5 µl of inoculum.

### MRI Acquisition

MRI acquisition was performed in a 7T BioSpec animal MRI from Bruker. This scanner was equipped with Avance III electronics and the 500V/300A B-GA12S2 gradient upgrade with a standard 40 mm quadrature volumetric transceiver. Animals were anesthetized with a 1% isoflurane/medical air mixture. A constant 37°C air flow was used to keep the animals warm. Structural imaging was obtained using the Bruker standard 3D-True Fast Imaging with Steady State Precession pulse sequence. To remove banding artifacts, a root-mean-square image of eight phase advance (angles of 0–315 degrees in increments of 45) acquisitions was obtained. Each TrueFISP phase angle acquisition was acquired as follows: slices oriented in the rostrocaudal axis, FOV of 36 × 36 × 36 mm with a matrix of 180 × 180 × 180, TE/TR of 2.5/5.0 ms, NEX of 2, flip angle of 30°, and a bandwidth of 50 kHz; no accelerations were used. The resulting image is an average of 16 acquisitions with an isotropic 200 μm resolution and was acquired in a total scanning time of 46 min.

### PET Imaging

PET acquisition was performed using a CTI Concorde R4 MicroPET for small animals. Animals were induced using 5% isoflurane in 1.0 L/min oxygen and then maintained with 2% isoflurane in medical air. A 9 min transmission scan was done using a rotating [57Co] point source (34). This was followed with a bolus injection of [^18^F]MK-6240 or [^18^F]NAV4694 in the tail vein. The emission scan began immediately after, which lasted for 60 minutes. The data from the dynamic scan were reframed into 27 sequential time frames of increasing durations (8× 30 sec, 6× 1 min, 5× 2 min, and 8× 5 min). MINC tools were used for image processing and analysis (www.bic.mni.mcgill.ca/ServicesSoftware), following the pipeline that was previously described by Parent et al. (35). Briefly, averaged PET images for each animal were coregistered to their respective structural MRI, which was then non-linearly transformed into the Waxholm Sprague Dawley Space Atlas (https://www.nitrc.org/projects/whs-sd-atlas). The binding potential (BP_ND_) maps were generated using the SRTM model, [^18^F]NAV-4694 BP_ND_ was calculated for each voxel using cerebellar gray matter as a reference region and for [^18^F]MK-6240 BP_ND_ was calculated for each voxel using the striatum (36). The parametric maps were blurred using a 1mm Full Width Half Maximum Gaussian kernel.

## Acknowledgements

We thank the radiochemistry team at the Montreal Neurological Institute for synthesizing [18F]MK-6240, especially Karen Ross, I-Huang Tsai, Robert Hopewell for MK-6240 productions. We are grateful to the Douglas Brain Bank and the families of donors for providing patient tissues. We also thank Dominique Mirault for coordinating the brain donations and Dr. Naguib Mechawar, the director of the Douglas Brain Bank. We thank the staff members from the Douglas Hospital animal facility for animal care and husbandry. We gratefully acknowledge the Cryo-Electron Microscopy Facility (CEMF) and Structural Biology Laboratory (SBL) at UT Southwestern Medical Center for training in data acquisition, including Drs. Daniel Stoddard, Jose Martinez, Raymond Welch from CEMF, and Drs. Zhe Chen, Yang Li, and Yan Han from SBL. We thank Rebecca Jackson and Phoebe Doss from the Electron Microscopy Core Facility at UTSW for training on Tecnai Spirit, and thank Michel Goedert for facilitating our introduction to Sofia Lovestam, who assisted with questions related to cryo-EM reconstruction using RELION 4.0. We extend thanks to Paul Seidler for discussions about the interpretation of the data presented in this manuscript.

## Author Contribution

The project was conceived and designed by PK, PRN, MD, and SS. Acquisition and histologic/diagnostic analysis of patient brain tissue were performed by MCG. JVA prepared cryo-EM grids and edited the manuscript. PK cut tissue and performed relative autoradiography, purified patient tissue samples, prepared grids, performed PET experiments and analyzed all data, performed cryo-EM image analysis, model building, and refinement, developed figures. JM helped with model refinement, helped movie generation with PK and SS, and edited the manuscript. GM and JPS oversaw all aspects related to radiopharmaceutical productions. JPS oversaw nuclear medicine procedures conducted in this manuscript. PRN oversaw all PET analytical pipelines and autoradiography experiments. MSS conducted the symmetry docking and SASA calculations under the supervision of BKS. PK and SS wrote the paper. MD, PRN, and SS edited the paper with input from all authors. SS supervised the project.

## Corresponding Author

Correspondence to Sarah Shahmoradian, Marc Diamond, and Pedro Rosa-Neto.

