## Extended Data for "Cryo-EM structure of Alzheimer’s disease tau filaments with PET ligand MK-6240"

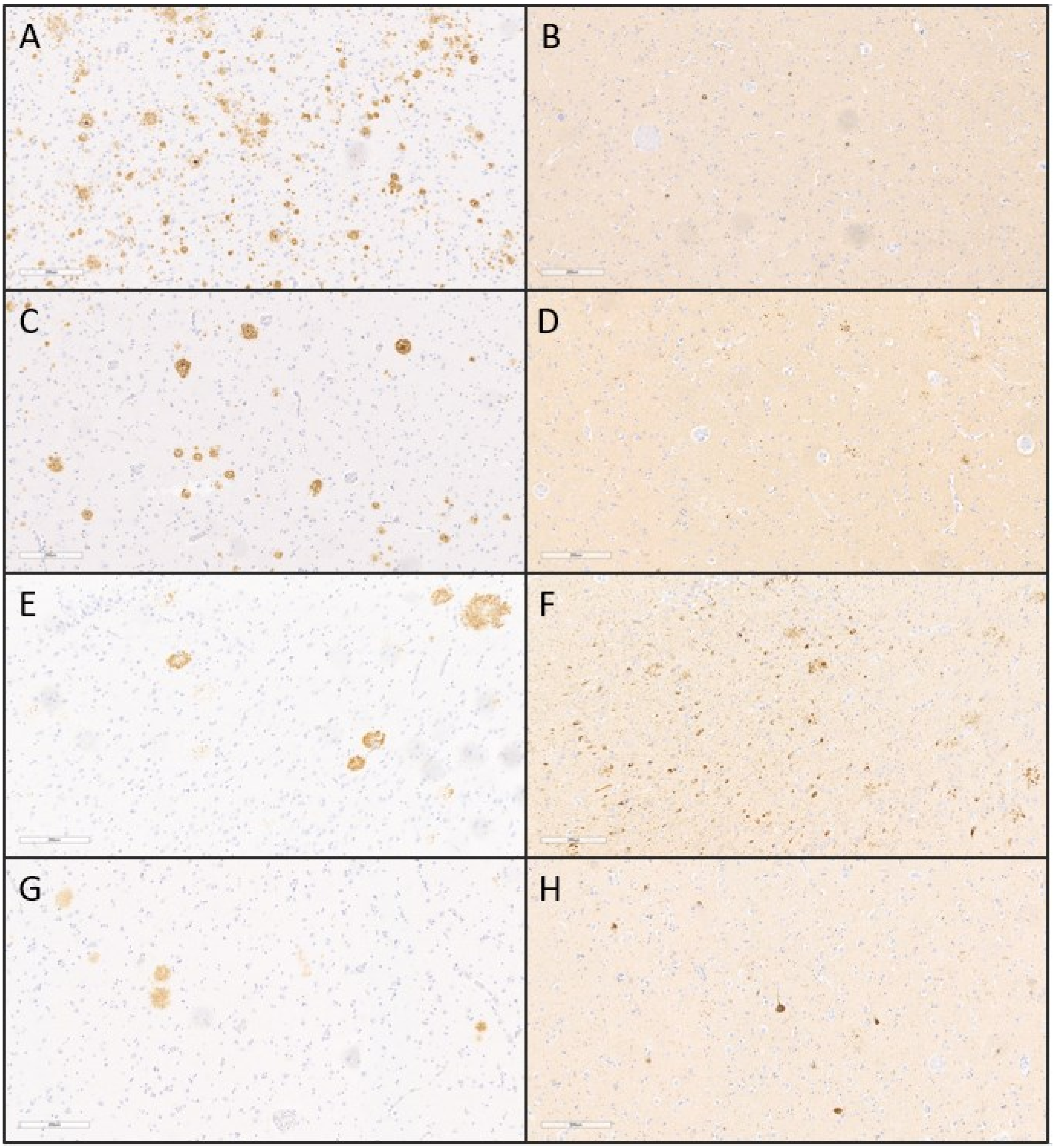


**Extended Data Figure 1: Tau and Amyloid Staining in Alzheimer's Disease Patient Utilized as Source of Tau Filaments.**

**(A)** Immunohistochemical staining of sections derived from the Parietal cortex (A and B), Medial temporal cortex (C and D), Hippocampus (E and F), and Amygdala (G and H) employing the 1-40/42 anti-amyloid antibody (A, C, E, and G) and AT8 (B, D, F, H). Nuclei were counterstained. Scale bars: 200 µm.

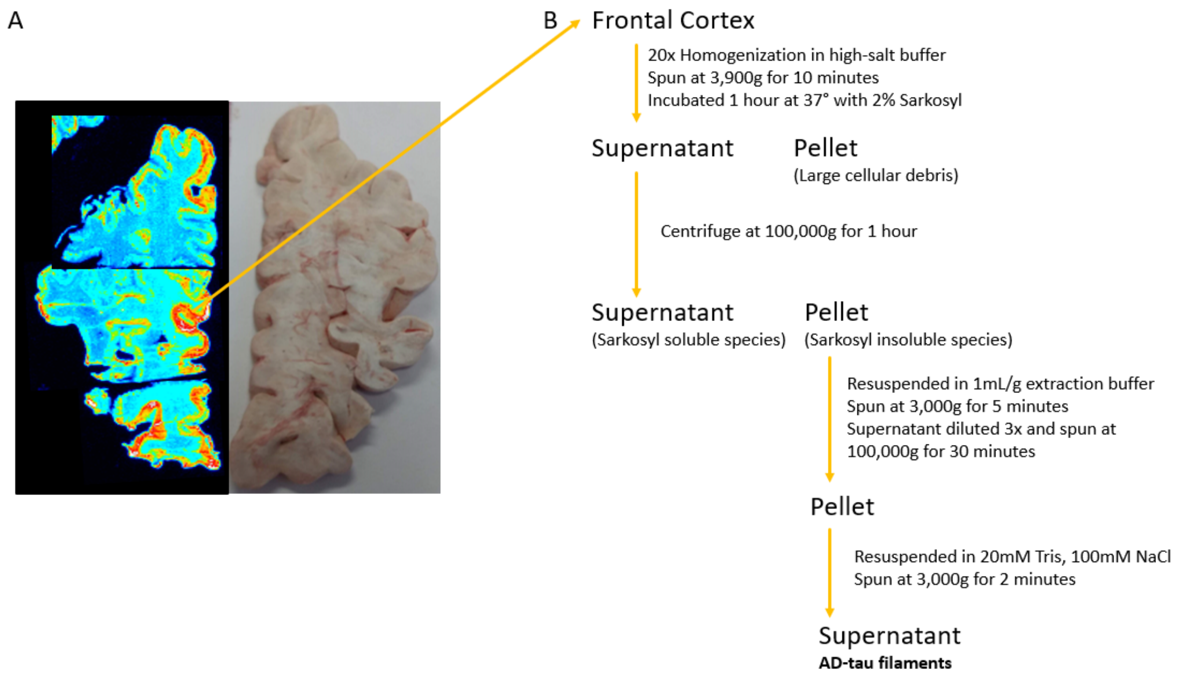


### Extended Data Figure 2: Schematic of Tissue-Guided Autoradiography and Homogenization.

**(A)** Autoradiograph of post-mortem brain tissue from a neuropathologically confirmed Alzheimer's disease (AD) patient, labeled with [18F] MK-6240. Color code denotes regions of robust [18F] MK-6240 binding in red and areas of low binding in blue. Grey matter regions with notable [18F] MK-6240 binding were isolated and homogenized for the extraction of sarkosyl-insoluble fractions. **(B)** Schematic representation outlining the sequential steps involved in the purification of AD-tau filaments.

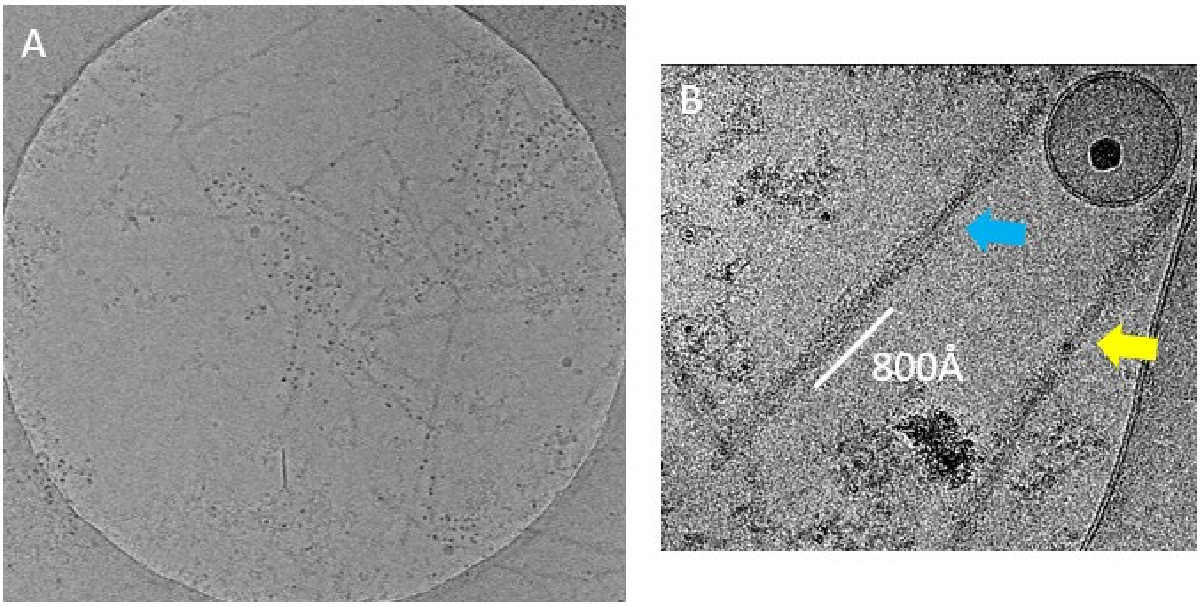


**Extended Data Figure 3: Micrograph Assessment for Data Collection Preparation.**
**(A)** Depiction of a representative view of the cryo-EM grid utilized for data collection. **(B)** Illustration of representative micrographs showcasing paired helical filaments (PHFs) in blue, accompanied by a lesser occurrence of straight filaments (SFs) in yellow. A cross-over distance of 800 Å was observed for PHFs, guiding the initial model generation.

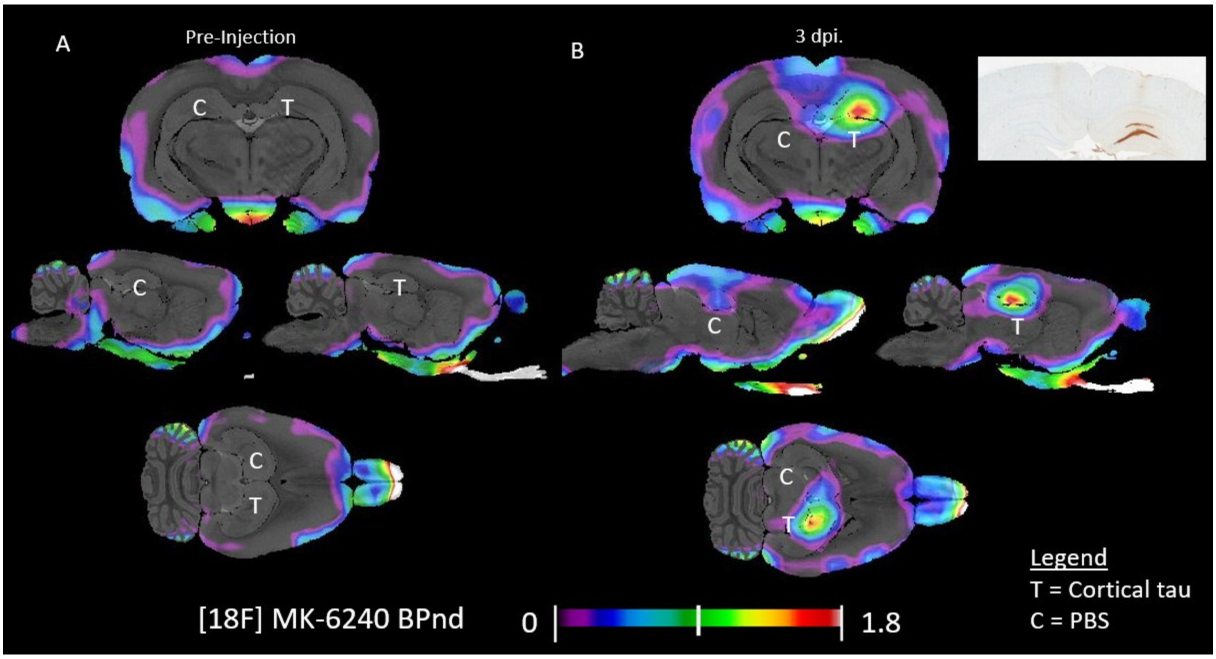


**Extended Data Figure 4: Pilot Inoculation Utilizing Sarkosyl-Insoluble Tau Filaments Extracted from Alzheimer's Disease Patient Brain.**

**(A)** Depicts the pre-injection [18F] MK-6240 BPND (n=1). **(B)** Displays the post-injection [18F] MK-6240 BPND, accompanied by post-mortem immunohistochemistry (IHC) evidencing AT8 positivity (n=1). The left hemisphere of the dorsal hippocampus received a control injection of PBS, while the right hemisphere was injected with sarkosyl-insoluble tau species extracted from human AD patient brain material.

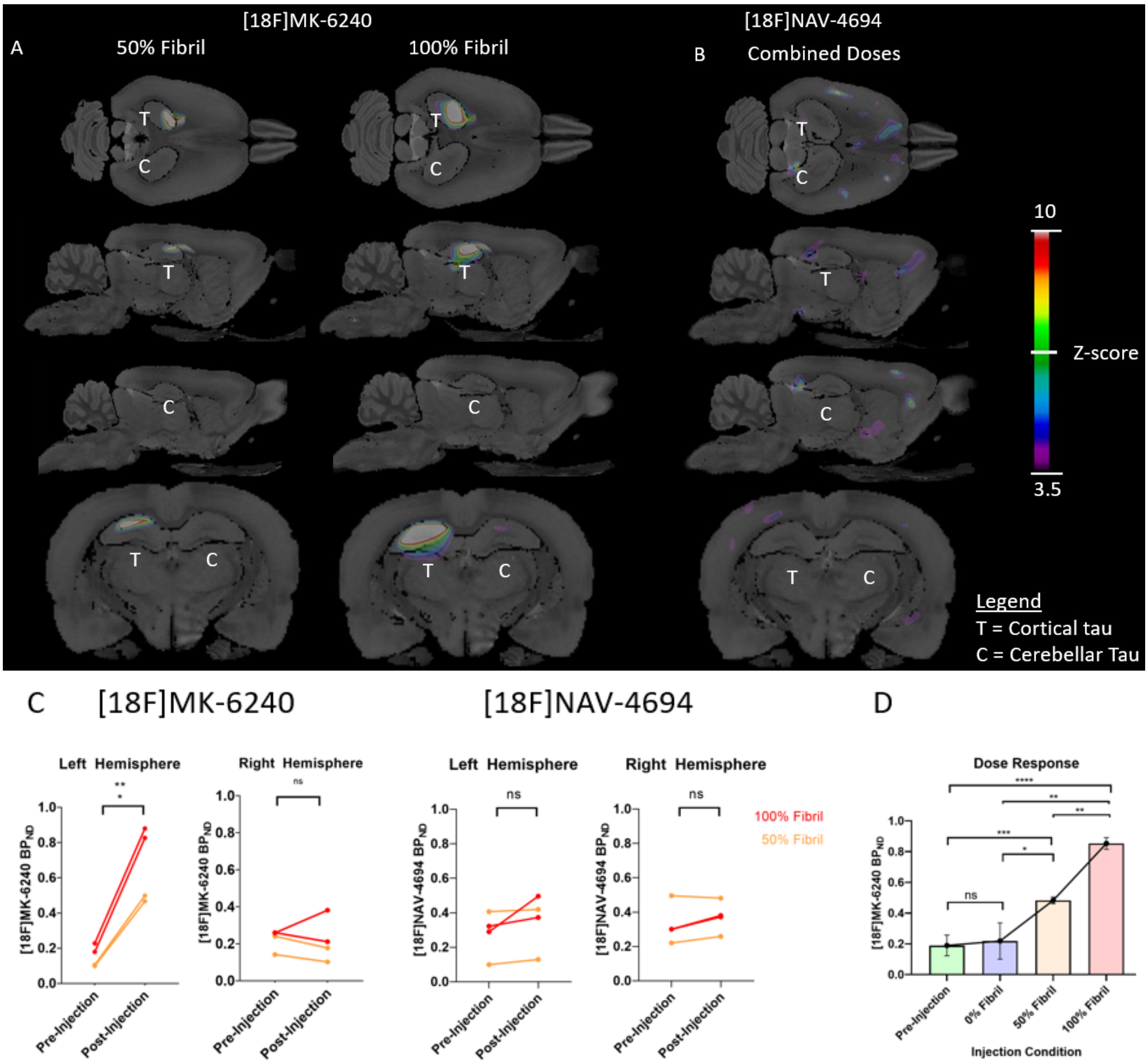


### Extended Data Figure 5: Sarkosyl-Insoluble Brain Material Quantification from Alzheimer's Disease Patient Brain in Living Rats using PET.

**(A)** Comparison of [18F] MK6240 z-score maps between baseline scans and post-injection scans in animals subjected to 50% Fibril dose (n=2) and 100% Fibril dose (n=2) AD tau fibril injections (Left Hemisphere), along with cerebellar cortex extraction control (Right Hemisphere). **(B)** [18F] NAV4694 z-score map comparison between baseline scans and post-injection scans in all animals injected with AD tau fibrils (n=4) (Left Hemisphere), juxtaposed with cerebellar cortex extraction control (Right Hemisphere). Overlaid on the Waxholm Space Atlas MRI. Brain maps were derived for [18F] MK6240 (tau-PET) and [18F] NAV-6494 (Aβ-PET) employing the formula [(Post-injection Average $BP_{ND}$) - (Baseline $BP_{ND}$ Average) / (Baseline $BP_{ND}$ Standard Deviation)]. This voxel-wise analysis was adjusted for multiple comparisons using the RFT method, yielding an adjusted threshold of $p < 0.05$. Post-injection $BP_{ND}$ Maps exhibit significant deviations from baseline, with z-scores ≥ 3.5. **(C/D)** Graphical representation displaying mean $BP_{ND}$ values originating from the dorsal hippocampi.

**Extended Data Table 1**
**Cryo-EM data collection, refinement, and validation statistics**

| **Data Collection** | **+MK-6240 AD PHF** | **-MK-6240 AD PHF** |
|---|---|---|
| Electron Microscope Type | Titan Krios | Titan Krios |
| Nominal Magnification | 105,000x | 105,000x |
| Voltage (kV) | 300 | 300 |
| Detector | Gatan K3 w/ Bioquantum Energy Filter | Gatan K3 w/ Bioquantum Energy Filter |
| Electron Exposure ($e^{-}$/ Å$^2$) | 62 | 60 |
| Defocus Range (μm) | -1.0 to -2.4 μm | -1.0 to -1.8 μm |
| Super-resolution Pixel Size (Å) | 0.415 | 0.415 |
| **Reconstruction** | | |
| Total number of micrographs | 8,037 | 12,477 |
| Number of usable micrographs | 3,976 | 3,842 |
| Particles after Extraction | 1,533,788 | 1,033,374 |
| Particles after 2D Classification | 408,700 | 413,684 |
| Particles after 3D Classification | 32,986 | 165,401 |
| Map Resolution (Å; FSC=0.143) | 2.31 | 3.08 |
| Helical Rise (Å) | 2.38 | 2.38 |
| Helical Twist (degrees) | 179.46 | 179.46 |
| Symmetry Imposed | $2_1$ | $2_1$ |
| **Refinement** | | |
| Initial model used (PDB code) | N/A | 5o3l |
| Model resolution FSC 0.5 (Å) | 3.10 | 2.92 |
| Map sharpening B factor (Å$^2$) | -33.04 | -65.83 |
| Model composition | | |
| Non-hydrogen atoms | 7236 | 7165 |
| Protein residues | 450 | 462 |
| Ligands | 6 | 0 |
| B Factors (Å$^2$) | | |
| Protein | 73.16 | 72.84 |
| Ligand | 0 | N/A |
| R.m.s. deviations | | |
| Bond lengths (Å) | 0.012 | 0.010 |
| Bond angles (degrees) | 1.946 | 1.931 |
| Validation | | |
| MolProbity score | 1.05 | 0.98 |
| Clashscore | 0.42 | 0 |
| Poor rotamers (%) | 0 | 0 |
| Ramachandran plot | | |
| Favored (%) | 94.06 | 92.22 |
| Allowed (%) | 5.94 | 7.78 |
| Disallowed (%) | 0 | 0 |

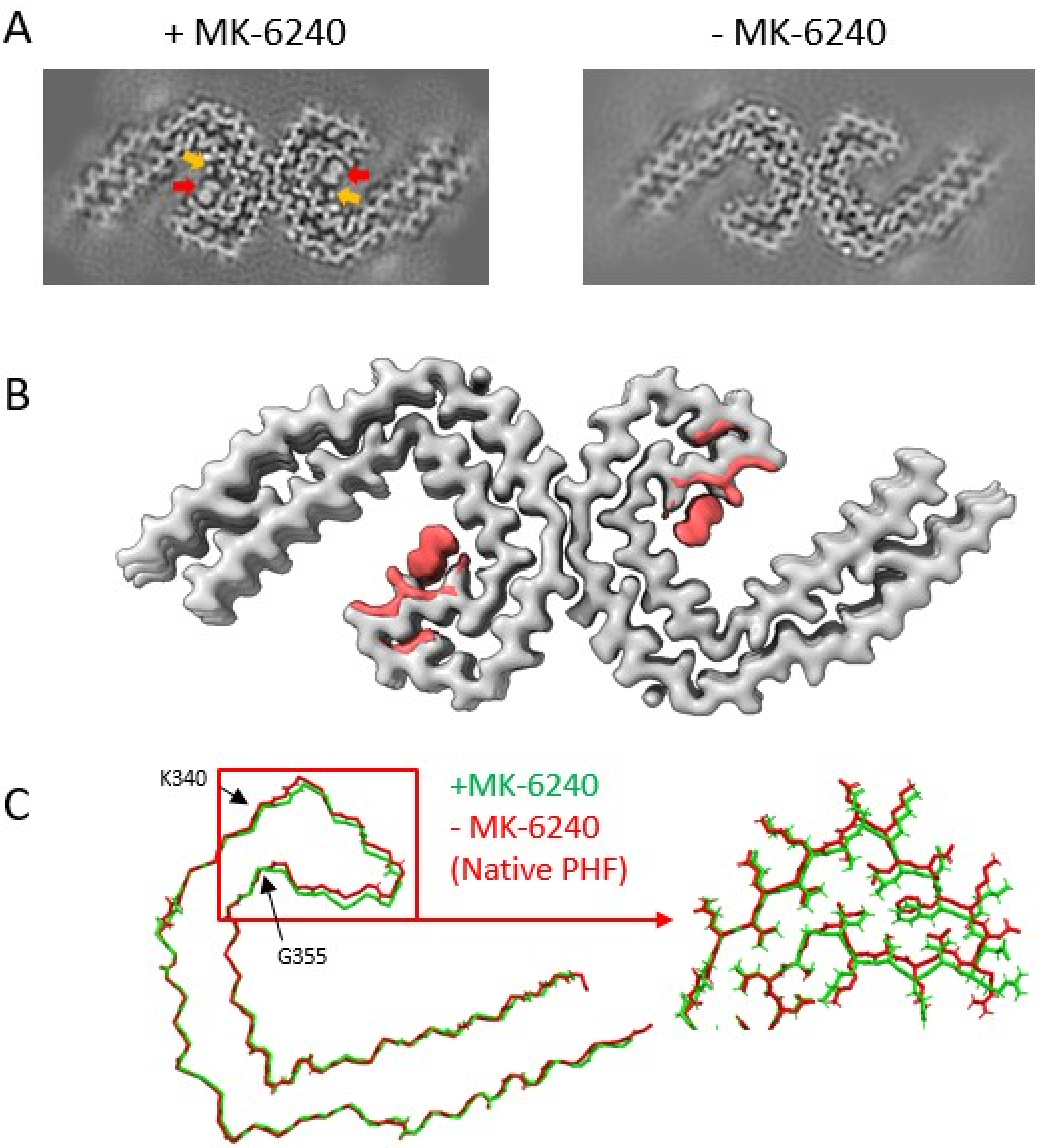


**Extended Data Figure 6: Structural Alterations and Maps Illustrating Mainchain and Sidechain Rearrangement upon MK-6240 Binding.**

**(A)** Projected cross-section of a single rung extracted from the final post-processed map. Cryo-EM map depicts the architecture of Alzheimer's disease (AD) paired helical filaments (PHFs) incubated with MK-6240 (left) (Unambiguous site is highlighted by a red arrow, second potential site is highlighted by an orange arrow), and without MK-6240 (DMSO vehicle control, right). Cross-section corresponds to six images, representing one rung of the filament. Sigma contrast value = 7.5. **(B)** Cryo-EM map of AD tau PHFs without MK-6240 (DMSO Control, grey) with the difference map derived from the MK-6240-bound structure (salmon), highlighting the most prominent differences, and emphasizing amino acids involved in sidechain rearrangements. **(C)** Overlaid atomic models displaying mainchains from native AD PHF (red) and +MK-6240 AD PHF (green). On the left, a mainchain translation originating at K340 and terminating at G355 is observable. The right section presents a magnified view from within the red rectangle on the left, illustrating distinct mainchain and sidechain configurations.

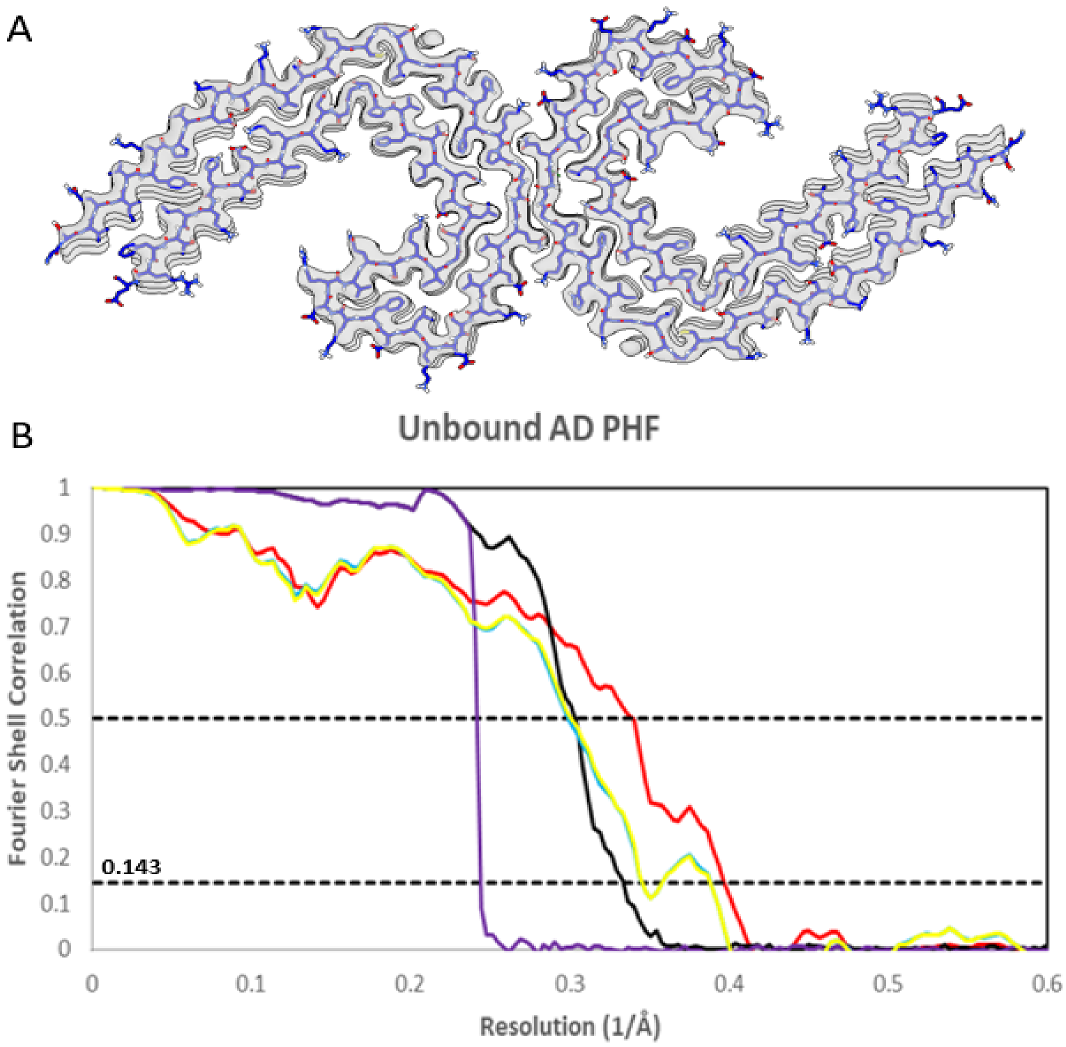


**Extended Data Figure 7: Cryo-EM Map and atomic model of unbound AD PHF structure.**

**(A)** Cryo-EM density (white) is juxtaposed with the atomic model of the unbound tau fold (blue). **(B)** Graph illustrating Fourier Shell Correlation (FSC) curves for the Cryo-EM maps of the AD-PHF with MK-6240 (Black). Additionally, FSC curves are presented for the refined atomic model against the post-processed map utilized for atomic model generation (Red), for the refined atomic model from one half-map to itself (Blue), for the refined atomic model from the first half-map to the second (Yellow), and for phase randomized curves of the two independently refined half-maps (Purple).

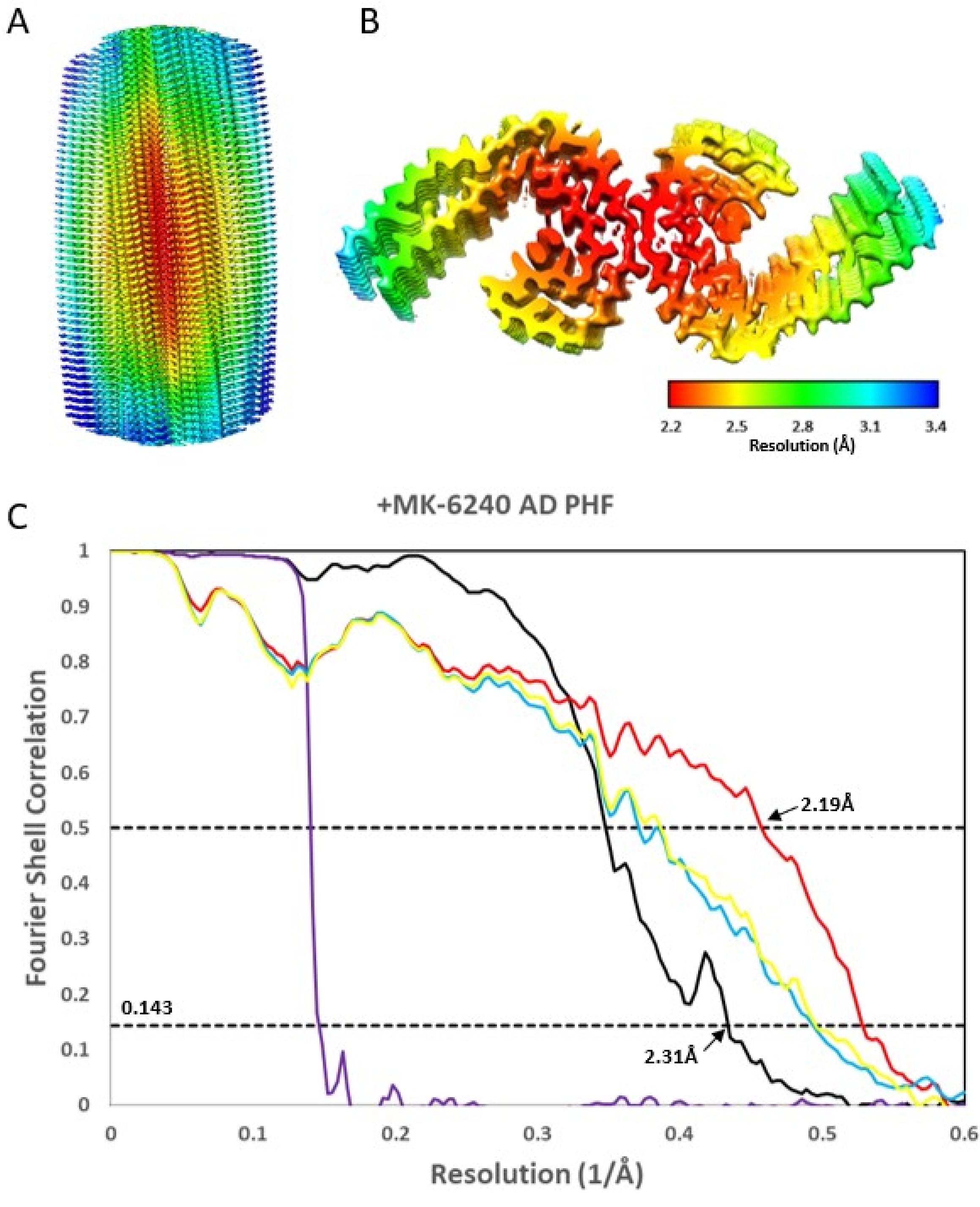


**Extended Data Figure 8: Cryo-EM Map with Local Resolution and Model Resolution Assessment.**

**(A/B)** Cryo-EM map depicting Alzheimer's disease (AD) paired helical filaments (PHF) with MK-6240, illustrating a side-view of the filament (A) and a cross-sectional view representing 20% of the filament (B). Local resolution is represented using color coding (in Å), where warmer colors indicate regions of high resolution and cooler colors indicate areas of lower resolution. **(C)** Graph illustrating Fourier Shell Correlation (FSC) curves for the Cryo-EM maps of the AD-PHF with MK-6240 (Black). Additionally, FSC curves are presented for the refined atomic model against the post-processed map utilized for atomic model generation (Red), for the refined atomic model from one half-map to itself (Blue), for the refined atomic model from the first half-map to the second (Yellow), and for phase randomized curves of the two independently refined half-maps (Purple).

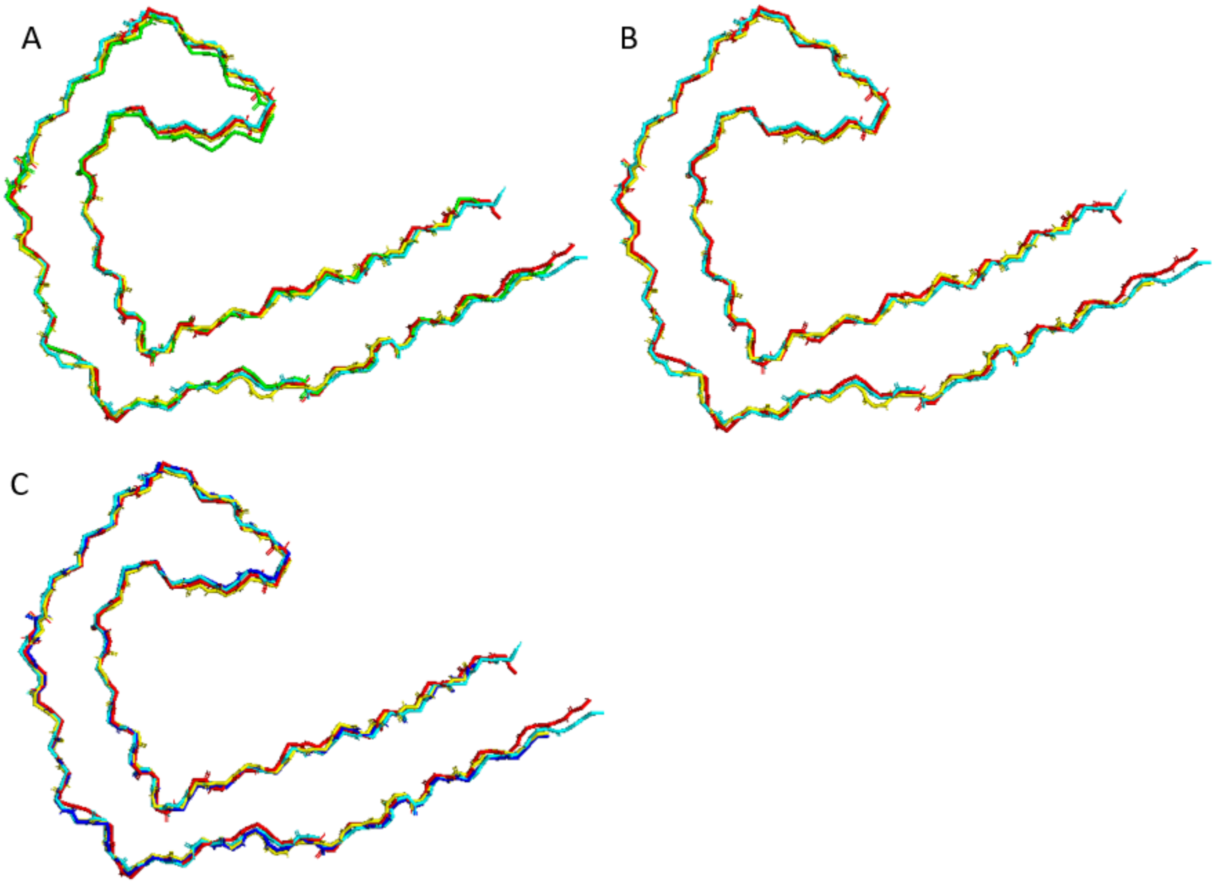


**Extended Data Figure 9: Atomic Model Comparison for RMSD Calculations**

**(A)** Overlaid atomic models from +MK-6240 AD PHF (green). unbound AD PHF (red), 503L (yellow), and 6HRE (baby blue). **(B)** Overlaid atomic models from unbound AD PHF (red), 503L (yellow), and 6HRE (baby blue). **(C)** Overlaid atomic models from +GTP-1 AD PHF (dark blue). unbound AD PHF (red), 503L (yellow), and 6HRE (baby blue).

**Surface Accessible Surface Area Calculations**

We translated and rotated a copy of the 3 Rung model refined from our cryo-EM map into a model containing 6 tau rungs because a single ligand interacts with 4 rungs. Running this calculation for the 3-rung model would have left some of the middle ligand hanging off the side of the modeled protein. Doing so undervalues the buried SASA, so we calculated these values for the 6-rung model.

We calculated the buried SASA for an MK-6240 monomer in the protein fibril using Chimera.

> The SASA of a single ligand in the fibril, is Z = 20328 $Å^2$
> The SASA of just the protein is P = 20293 $Å^2$
> Then the SASA of MK-6240 buried in the protein is (X + P - Z) / 2 = 208 $Å^2$ or 46% of the MK-6240 monomer surface area.

Next, we calculated the buried SASA for an MK-6240 monomer in a ligand stack.

> The SASA of a ligand dimer is D = 657 $Å^2$
> The SASA of a ligand trimer is T = 865 $Å^2$
> Then the buried SASA in just the ligand stack is 2X - D = 244 Å2, or X - (T - D) = 243 $Å^2$ or 54% of the MK-6240 monomer surface area.

Lastly, to be consistent with the calculations from the literature (30), we calculated the buried SASA for an MK-6240 monomer in a stack of ligand in the protein fibril.

> The SASA of a protein fibril with 3 bound ligands in a stack (S), of a fibril with 2 bound ligands separated by a site between them (N), and of a single ligand monomer out in space (X). Like the calculation with a single ligand in the fibril, the buried SASA at the ligand-protein interface and ligand-ligand interface is:
>
> (X + N - S) / 2
> X = 451 $Å^2$, N = 20373 $Å^2$, and S = 20201 $Å^2$.
> Thus, the buried SASA is 311 $Å^2$, or 69% of MK-6240 monomer surface area.
